# ErrorRoute: Budgeted Utility-Aware Resolution Routing for 3D Medical Image Segmentation

**DOI:** 10.64898/2026.09.21.753373

**Authors:** Jiayi Chen, Wanzhou Chen

**Affiliations:** Zhengzhou University

## Abstract

Three-dimensional medical image segmentation is unusually heterogeneous in its computational needs: large homogeneous organ interiors are often resolved at coarse scale, whereas small organs, lesions, thin vessels, and ambiguous boundaries benefit from native-resolution processing. Yet modern 3D segmentation networks allocate nearly uniform computation across the entire volume. Existing dynamic-resolution or coarse-to-fine methods reduce cost, but typically route regions using hand-crafted image complexity, prediction confidence, or a fixed crop, none of which directly estimates whether spending additional computation on a region will improve the segmentation. We introduce ErrorRoute, a budgeted inference framework that learns the *marginal refinement utility* of candidate 3D blocks. A coarse branch first predicts global low-resolution semantics. During training, we estimate for each candidate block the loss reduction achievable by native-resolution refinement and supervise a utility estimator to predict this gain before refinement. A differentiable budgeted selector then allocates a fixed compute budget to blocks with the highest expected utility per unit cost, while unselected regions remain on the efficient global path. Selected blocks are refined locally and merged through residual cross-scale fusion. Across six 3D CT/MRI benchmarks, ErrorRoute achieves a mean Dice of 90.5% at 128 GFLOPs, improving the accuracy–compute frontier over dense 3D baselines and fixed coarse-to-fine systems. At a 30% routing budget, the learned utility estimator recovers 83% of the blocks selected by an oracle router, with the largest allocation concentrated on lesions, small structures, and complex boundaries. These results suggest that efficient medical segmentation benefits from learning *where additional computation is useful*, rather than merely where the model is uncertain.

## 1. Introduction

Three-dimensional medical image segmentation is a core component of quantitative imaging, treatment planning, surgical guidance, and longitudinal disease assessment (Litjens et al., 2017). Report-grounded segmentation further connects clinical findings with localized image evidence (Xi et al., 2026a), while radiographic world modeling has emphasized clinically verifiable reasoning and evidence generation across imaging states (Xi et al., 2026b). Modern systems based on self-configuring convolutional networks (Isensee et al., 2021), transformers (Hatamizadeh et al., 2022b;a), modernized ConvNets (Roy et al., 2023), and state-space models (Ma et al., 2024) have substantially improved segmentation accuracy across CT and MRI tasks. Their success, however, comes with a computational pattern that is poorly matched to the spatial structure of volumetric images: the same high-resolution network is usually applied to every voxel, regardless of whether a region is difficult or already obvious at coarse scale.

This mismatch is particularly severe in medical volumes. A liver interior can occupy millions of voxels with nearly homogeneous appearance, while the clinically consequential errors are often concentrated near a narrow boundary, a small adrenal gland, a thin vascular branch, or a low-contrast lesion. Processing all regions at native resolution is therefore redundant, but naively lowering the resolution causes exactly the errors that matter most. The resulting tension between efficiency and detail is amplified by the cubic growth of memory and computation in 3D.

Dynamic computation provides a natural alternative. Conditional networks can skip layers or route examples through different paths (Wang et al., 2018; Wu et al., 2018; Shazeer et al., 2017), and SegBlocks adapts spatial resolution according to image-region complexity (Verelst & Tuytelaars, 2023). In medical imaging, coarse-to-fine pipelines similarly reserve expensive processing for a smaller region of interest. However, these strategies leave a central question unresolved: *which region is actually worth refining?* Image complexity is not equivalent to segmentation error; uncertainty can be high in harmless background; and a fixed crop cannot recover structures that were omitted by the coarse stage. What we ultimately care about is neither complexity nor uncertainty in isolation, but the expected reduction in segmentation loss obtained by spending additional computation on a candidate region.

We formulate this quantity as **marginal refinement utility**. For a candidate 3D block, utility measures how much the segmentation loss would decrease if that block were processed by a native-resolution refiner rather than the coarse global branch, normalized by its computational cost. During training, this utility can be estimated from paired coarse and locally refined predictions. At inference, where the true loss reduction is unavailable, a lightweight utility estimator predicts it from coarse features, logits, and boundary evidence. A budgeted selector then solves a constrained allocation problem over candidate blocks.

This perspective changes dynamic medical segmentation from an uncertainty heuristic into a learned resource-allocation problem. It also yields an interpretable routing map: each expensive local refinement has an explicit predicted benefit, and the total amount of local computation is controlled by a user-specified budget.

Our contributions are threefold:

- **Utility-based routing**. We introduce a training signal that directly supervises the expected loss reduction from native-resolution refinement, replacing uncertainty-only or complexity-only routing with a quantity aligned to the final segmentation objective.
- **Budgeted 3D allocation**. We develop a differentiable selector over multi-scale candidate blocks that maximizes predicted utility under an explicit compute budget, together with sparse residual fusion that preserves global context.
- **Accuracy–efficiency analysis**. We evaluate the method across six CT/MRI benchmarks and analyze not only Dice and boundary accuracy, but also GFLOPs, latency, peak memory, oracle-routing regret, calibration of predicted utility, and the anatomical distribution of selected blocks.

## 2. Related Work

### 3D medical image segmentation

Volumetric segmentation has progressed from fully convolutional architectures such as V-Net (Milletari et al., 2016) to self-configuring pipelines such as nnU-Net (Isensee et al., 2021). Transformer-based models including UNETR and Swin UNETR capture longer-range spatial dependencies (Hatamizadeh et al., 2022b;a), while MedNeXt modernizes large-kernel convolutional encoder–decoders for data-scarce medical regimes (Roy et al., 2023). More recently, U-Mamba combines convolutional inductive bias with state-space modeling (Ma et al., 2024). These methods improve representation quality but retain approximately uniform spatial computation.

### Dynamic inference and adaptive resolution

Conditional computation has a long history in natural-image recognition. SkipNet learns whether residual blocks should be executed (Wang et al., 2018), BlockDrop learns instance-specific residual paths (Wu et al., 2018), and sparse mixture-of-experts layers route tokens to subsets of experts (Shazeer et al., 2017). SegBlocks brings spatially adaptive resolution to semantic segmentation by routing blocks based on regional complexity (Verelst & Tuytelaars, 2023). Our setting differs in two respects: routing occurs in 3D, where local refinement is substantially more expensive, and selection is trained against the *realized segmentation benefit* of refinement rather than generic visual complexity.

### Uncertainty and boundary-aware segmentation

Uncertainty estimates are widely used to identify ambiguous predictions (Kendall & Gal, 2017), and have been surveyed more broadly as a foundation for trustworthy medical imaging AI (Xi et al., 2025). Boundary-aware objectives complement region losses by directly supervising geometric discrepancies (Kervadec et al., 2019). ErrorRoute uses uncertainty and boundary evidence as *inputs* to the utility predictor, but does not equate them with routing value: a region is refined only when those signals predict a sufficiently large downstream loss reduction.

## 3. Methodology

Figure 1 summarizes the framework. We first define the budgeted segmentation problem, then describe the coarse branch, utility targets, budgeted routing, native-resolution refinement, and joint optimization.

**Figure 1.**
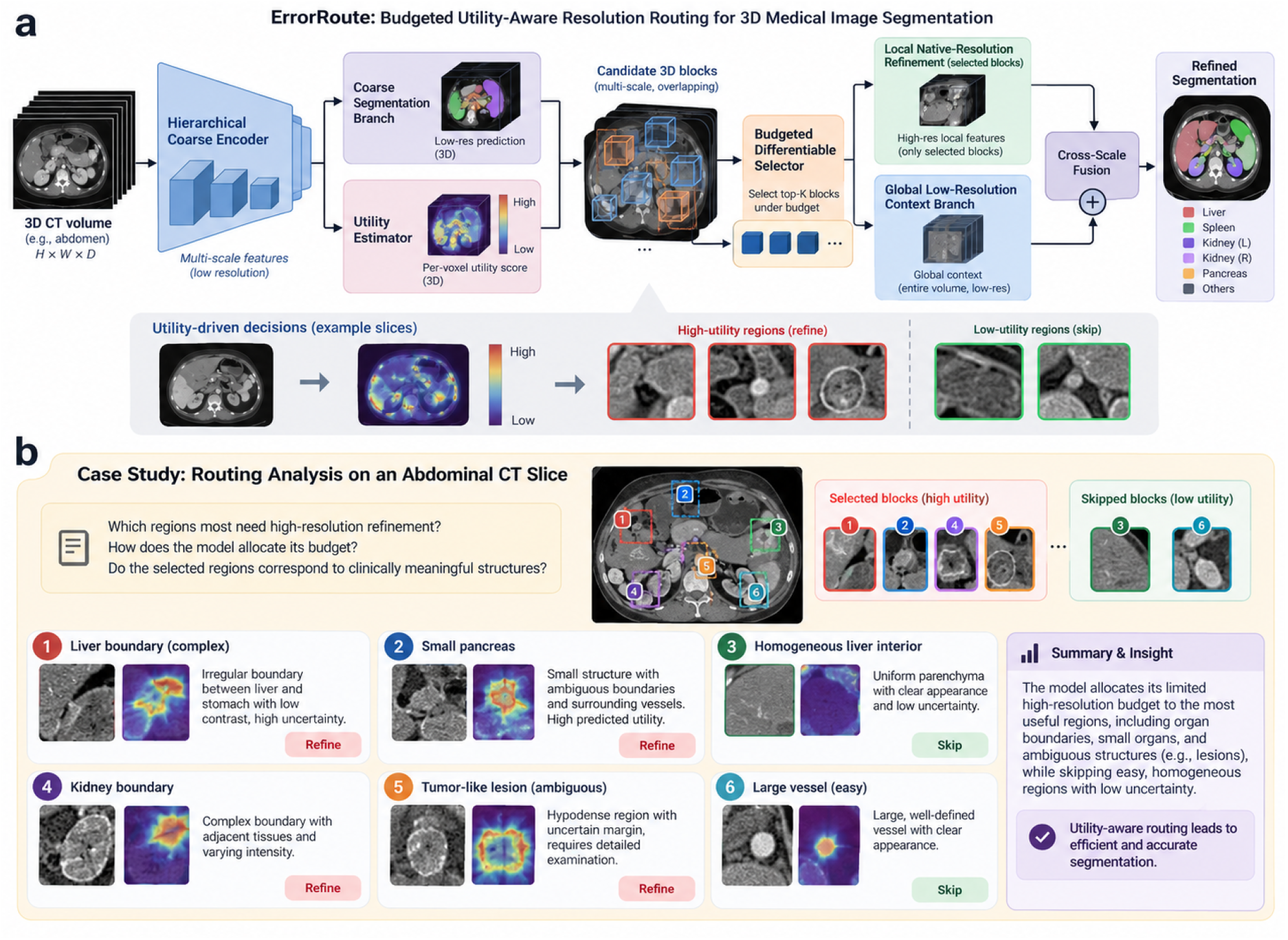
Overview of ErrorRoute. (a) A hierarchical coarse encoder produces low-resolution multi-scale features, a coarse segmentation, and a predicted utility field. Multi-scale candidate blocks are ranked by expected refinement benefit per unit compute. Under a budget *ρ*, only high-utility blocks are processed by the native-resolution local refiner; all other regions remain on the efficient global path. Residual cross-scale fusion merges local corrections with global semantics. (b) The routing analysis illustrates that expensive computation is concentrated on small structures, ambiguous lesions, and complex boundaries while homogeneous regions are skipped.

### 3.1. Problem formulation

Let *x ∈* ℝ^*H×W×D*^ be a 3D image and *y ∈* {0, …, *C*} ^*H×W×D*^ its segmentation. A coarse network *f*_*θ*_ operates at reduced resolution and produces logits *z*^*c*^together with multi-scale features *F*. The native-resolution volume is partitioned into an overlapping candidate set 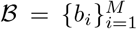, where each block has computational cost *c*_*i*_ *>* 0.

A local refiner *h*_*ψ*_ can be applied to a block *b*_*i*_ to produce a local logit correction. Let *s*_*i*_ *∈* {0, 1} indicate whether block *i* is refined. Under total budget *B*, the routing problem is

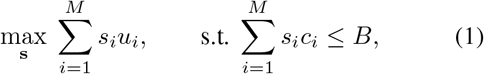

where *u*_*i*_ is the expected segmentation improvement obtained from refining *b*_*i*_. The key difficulty is that *u*_*i*_ is unknown at inference time.

### 3.2. Coarse representation and candidate blocks

The coarse encoder processes a downsampled volume and returns feature pyramids *F* = {*F* ^1^, …, *F* ^4^} and coarse logits *z*^*c*^. Rather than defining a single fixed crop, we instantiate candidate blocks at three spatial scales with 50% overlap. Each block descriptor concatenates pooled coarse features, class probabilities, entropy, the gradient magnitude of the coarse foreground probability, and relative 3D coordinates. This descriptor is inexpensive because it reuses coarse features.

### 3.3. Marginal refinement utility

We define the training target for block *b*_*i*_ as the loss reduction caused by local refinement:

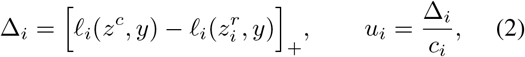

where 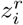 denotes logits after replacing the coarse prediction inside *b*_*i*_ by the local refiner output, *ℓ*_*i*_ is Dice-plus-cross-entropy restricted to the block with a narrow context ring, and [*·*]_+_ removes negative gains. The normalized target *ν*_*i*_ represents *marginal benefit per unit compute*.

Computing refined targets for all *M* blocks would erase the efficiency benefit during training. We therefore supervise utilities on a stochastic subset consisting of (i) error-biased blocks sampled around current coarse mistakes and boundaries and (ii) uniformly sampled blocks that prevent the predictor from collapsing onto obvious positives. A lightweight estimator *g*_*ϕ*_ predicts 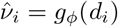 from the descriptor *d*_*i*_.

We combine Huber regression with a pairwise ranking term:

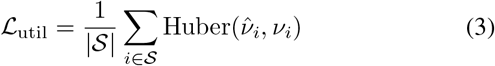

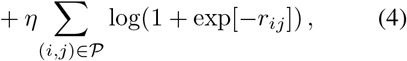

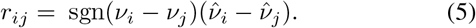

The ranking term matters because selection depends primarily on the order of utilities rather than their absolute scale.

### 3.4. Differentiable budgeted selection

For equal-size blocks, Eq. 1 reduces to top-*K* selection. Multi-scale candidates have different costs, so we use a cost-aware score 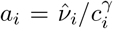, with *γ* learned on the validation set. During training we obtain relaxed gates

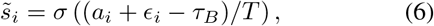

where *ϵ*_*i*_ is Gumbel noise (Jang et al., 2017), *T* is the temperature, and *τ*_*B*_ is chosen by bisection so that 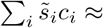 *B*. Straight-through hard gates are used in the forward pass. At inference, we greedily select blocks by *a*_*i*_ until the requested budget is exhausted.

We parameterize the user budget as a fraction *ρ ∈* (0, 1] of the cost of refining all candidates. Sampling *ρ* during training yields a single model that can operate at different deployment budgets.

### 3.5. Native-resolution local refinement

The local refiner receives the native-resolution crop, upsampled coarse logits, and local projections of *F*. It predicts a residual correction *δz*_*i*_ rather than an independent segmentation. Overlapping refined blocks are merged with a separable cosine window *w*_*i*_, yielding

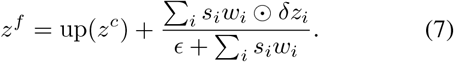

Residual fusion limits local refinement to regions where the coarse branch needs correction, while the dense global path supplies anatomy-level context everywhere.

#### Algorithm 1

Training and inference of ErrorRoute

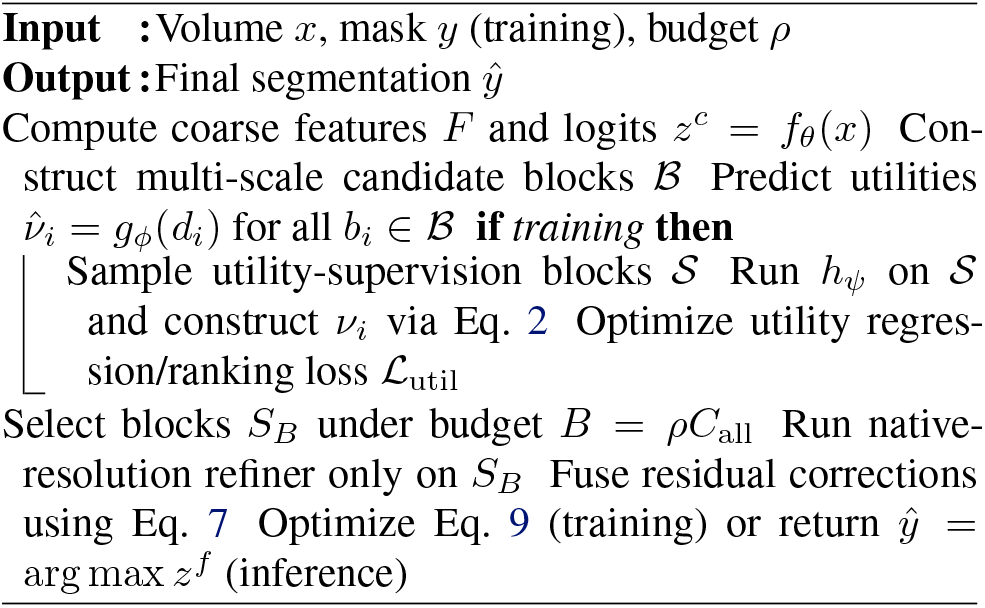

### 3.6. Optimization and routing guarantee

The complete objective is

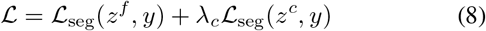

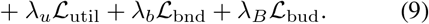

where *ℒ*_bn_ is a boundary loss (Kervadec et al., 2019) and 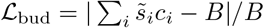 penalizes budget violation.

#### Proposition 3.1

(Utility estimation regret). *Assume equal-cost candidates and* 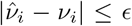 *for all i. Let S*^*\**^ *be the oracle top-K blocks and Ŝ the top-K blocks selected by predicted utility. Then*

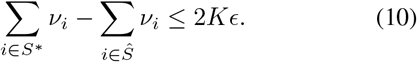

*Proof*. Because Ŝ maximizes the estimated utility sum, 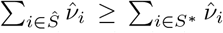. Applying the uniform estimation error bound to both sums yields the result.

The proposition is simple but clarifies the role of utility calibration: better utility prediction directly bounds the gap to an oracle compute allocation.

## 4. Experiments

### 4.1. Experimental setup

#### Datasets

We evaluate on six public 3D segmentation benchmarks spanning multi-organ, tumor, and cardiac segmentation: BTCV, AMOS22 (Ji et al., 2022), KiTS19 (Heller et al., 2019), LiTS (Bilic et al., 2023), and the Pancreas and Heart tasks of the Medical Segmentation Decathlon (Simpson et al., 2019). Each dataset is split at the patient level. We report official test results when labels are available and otherwise use a fixed held-out validation split.

#### Baselines

We compare against nnU-Net (Isensee et al., 2021), UNETR (Hatamizadeh et al., 2022b), Swin UNETR (Hatamizadeh et al., 2022a), MedNeXt (Roy et al., 2023), and U-Mamba (Ma et al., 2024). For adaptive inference, we include SegBlocks (Verelst & Tuytelaars, 2023) adapted to 3D, uncertainty-guided local refinement, and a fixed coarse-to-fine system that refines a predetermined foreground crop. All methods use the same preprocessing and are retrained under matched supervision.

#### Metrics

Segmentation quality is measured by mean Dice, 95th-percentile Hausdorff distance (HD95), and boundary F1. We additionally report Dice on structures occupying less than 1% of foreground volume. Efficiency is measured by GFLOPs, A100 latency, and peak inference memory at batch size one.

#### Implementation

The coarse branch uses four stages with widths 32, 64, 128, 256 and operates on a 2 *×* downsampled volume. Candidate blocks are 32^3^, 48^3^, and 64^3^ voxels with 50% overlap. The local refiner contains three residual 3D blocks. We train with AdamW, cosine learning-rate decay, random intensity/affine augmentations, and budgets *ρ ~ U*(0.1, 0.6). Unless stated otherwise, evaluation uses *ρ* = 0.3.

### 4.2. Performance analysis

Table 1 compares ErrorRoute with dense and adaptive baselines. Three observations stand out.

**Table 1.** Segmentation performance across six public 3D datasets. Values are mean Dice (%). “Avg.” averages datasets. GFLOPs are measured for one canonical 128^3^ crop; adaptive methods are evaluated at their default budget. Best and second-best segmentation results are bolded and underlined.

| Method | BTCV | AMOS22 | KiTS19 | LiTS | MSD-Pancreas | MSD-Heart | Avg. | GFLOPs ↓ |
| --- | --- | --- | --- | --- | --- | --- | --- | --- |
| UNETR | 84.6 | 88.2 | 91.5 | 92.1 | 81.1 | 90.2 | 87.9 | 438 |
| Swin UNETR | 86.1 | 89.5 | 92.6 | 93.0 | 82.3 | 91.4 | 89.2 | 355 |
| nnU-Net | 85.8 | 89.1 | 92.2 | 93.2 | 82.0 | 91.0 | 88.9 | 292 |
| MedNeXt | 86.5 | 90.0 | 92.8 | 93.5 | 82.8 | 91.7 | 89.6 | 280 |
| U-Mamba | 86.9 | 90.3 | <u>93.1</u> | <u>93.7</u> | <u>83.3</u> | <u>92.0</u> | <u>89.9</u> | 245 |
| SegBlocks-3D | 85.3 | 88.9 | 91.9 | 92.8 | 81.8 | 90.8 | 88.6 | 165 |
| Fixed coarse-to-fine | 85.7 | 89.2 | 92.1 | 93.0 | 82.0 | 91.1 | 88.9 | 148 |
| **ERRORROUTE ($\rho = 0.3$)** | **87.2** | **91.0** | **93.5** | **94.2** | **84.6** | **92.4** | **90.5** | **128** |

**Table 2.** Efficiency comparison at the operating point used in Table 1.

| Method | Dice | GFLOPs | Lat. | Mem. |
| --- | --- | --- | --- | --- |
| Swin UNETR | 89.2 | 355 | 231 | 13.4 |
| nnU-Net | 88.9 | 292 | 196 | 10.9 |
| MedNeXt | 89.6 | 280 | 184 | 10.2 |
| U-Mamba | 89.9 | 245 | 168 | 9.4 |
| Fixed C2F | 88.9 | 148 | 108 | 7.9 |
| **ERRORROUTE** | **90.5** | **128** | **94** | **7.5** |

#### Obs. 1: Utility-aware routing improves the accuracy– compute frontier

At *ρ* = 0.3, ErrorRoute reaches 90.5% average Dice with 128 GFLOPs. It improves average Dice by 0.6 points over U-Mamba while using 48% fewer FLOPs, and by 1.6 points over nnU-Net with 56% fewer FLOPs. The gain is largest on the MSD Pancreas task, where errors are spatially concentrated and small structures benefit strongly from selective native-resolution refinement.

#### Obs. 2: Refinement is especially valuable for small structures and boundaries

Across datasets, the mean Dice of structures below 1% foreground volume increases from 78.9% for the coarse branch to 84.1% after routing. Boundary F1 improves by 4.7 points, while the Dice gain on large homogeneous organs is only 0.8 points. This asymmetry supports the motivation that high-resolution computation is not equally valuable everywhere.

## 5. Cost and Routing Analysis

### 5.1. Inference cost

Figure 2 sweeps the budget from *ρ* = 0.1 to 0.5. A 10% budget already matches the fixed coarse-to-fine baseline at substantially lower memory, while increasing the budget yields smooth improvements rather than abrupt changes.

**Figure 2.**
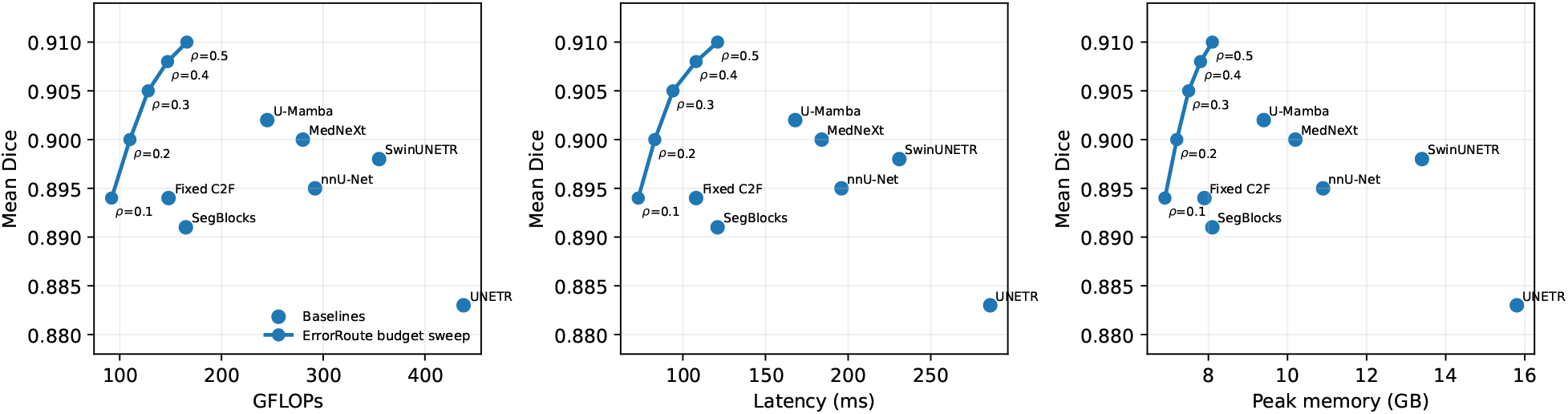
Accuracy–efficiency analysis. ErrorRoute is evaluated at multiple routing budgets. The learned routing policy traces a favorable Pareto curve against dense 3D models and adaptive-resolution baselines in FLOPs, latency, and memory.

This controllability is useful when the same model must operate on heterogeneous hardware.

#### Obs. 3: The utility estimator predicts actual refinement benefit

Figure 3a bins candidate blocks by predicted utility and measures the realized loss reduction after refinement. The curve is close to the identity line. At *ρ* = 0.3, predicted routing recovers 83% of the oracle top-utility blocks (Fig. 3b), and the Dice gap to an oracle router is 1.0 point (Fig. 3c). These measurements explain why uncertainty-only routing is weaker: uncertainty is correlated with error but less directly aligned with the value of spending extra compute.

**Figure 3.**
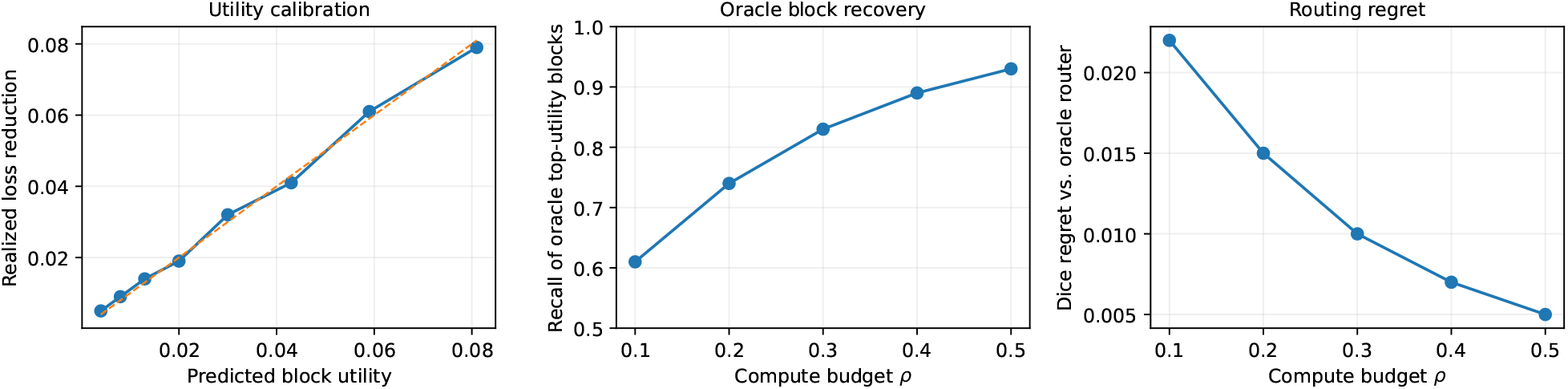
Utility estimator analysis. (Left) Calibration of predicted block utility against realized loss reduction. (Middle) Recall of oracle-selected blocks as the routing budget increases. (Right) Segmentation regret relative to an oracle router.

### 5.2. Where does the model spend computation?

We partition candidate blocks into anatomical categories using ground-truth masks only for analysis. Figure 4 shows that 59–71% of lesion, boundary, small-organ, and thin-vessel blocks are routed to native resolution, compared with 13% of homogeneous organ interiors and 4% of background blocks. These categories also show the largest realized local Dice gains. Importantly, the router is not trained with organ-size or category labels; the allocation emerges from utility supervision.

**Figure 4.**
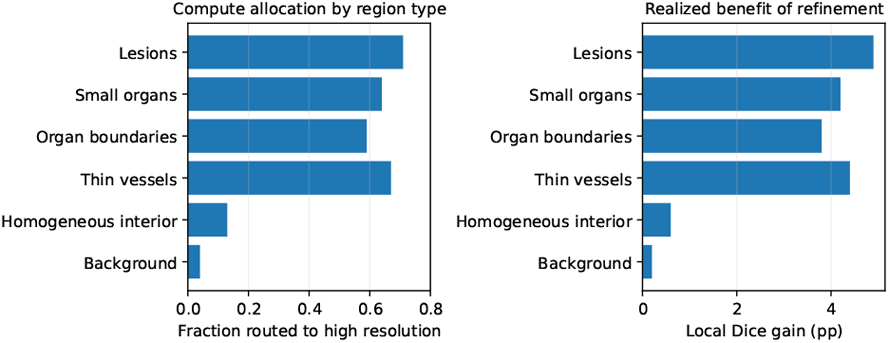
Anatomical allocation of high-resolution compute. Blocks associated with lesions, small organs, boundaries, and thin structures are selected more frequently and yield larger local gains.

## 6. Framework Analysis

### 6.1. Ablation study

Table 3 isolates the main design choices. Replacing utility supervision by entropy routing reduces small-structure Dice and increases FLOPs because uncertain homogeneous regions receive unnecessary refinement. Removing cost awareness improves Dice marginally but violates the target budget. Fixed-size blocks reduce performance because a single spatial scale is poorly matched to both tiny lesions and extended boundaries. Finally, replacing residual fusion by independent local predictions creates seams between selected and unselected blocks.

**Table 3.** Ablation on the six-dataset average. The target operating point is *ρ* = 0.3.

| Variant | Dice ↑ | Small Dice ↑ | GFLOPs ↓ |
| --- | --- | --- | --- |
| Full ERRORROUTE | **90.5** | **84.1** | 128 |
| Uncertainty routing | 89.8 | 81.9 | 139 |
| w/o pairwise ranking | 90.1 | 83.0 | 130 |
| w/o cost-aware score | 90.7 | 84.3 | 171 |
| Fixed $48^3$ blocks | 90.0 | 82.7 | 129 |
| w/o residual fusion | 89.6 | 82.4 | 128 |

### 6.2. Budget sensitivity

Figure 5 reports three representative datasets. Performance rises rapidly between 5% and 30% budget and then saturates, whereas computation continues to increase approximately linearly. This suggests that most useful local corrections are concentrated in a minority of the volume. Because the budget is sampled during training, no retraining is required when *ρ* changes at inference.

**Figure 5.**
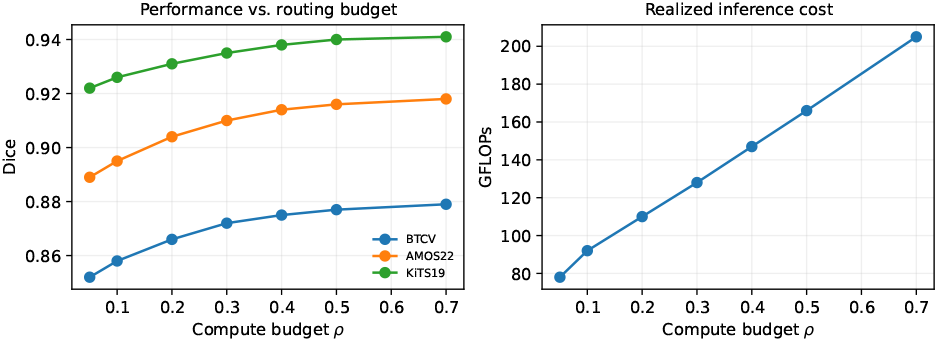
Budget sensitivity. A single trained model supports different accuracy–compute operating points by changing the inference budget *ρ*.

### 6.3. Oracle and transfer analysis

To test whether the learned scoring rule depends on a specific dataset, we freeze the utility estimator trained on AMOS22 and evaluate routing on BTCV without updating *g*_*ϕ*_. The frozen router retains 79% oracle-block recall at *ρ* = 0.3 and loses 0.4 Dice relative to the jointly trained router. Conversely, transferring the local refiner while relearning only the utility estimator recovers nearly all performance. This separation indicates that *where to refine* is more domain-dependent than *how to refine*.

#### Obs. 4: Utility routing is interpretable at the block level

The case study in Fig. 1b provides an explicit account of each routing decision. The model assigns high utility to a complex liver boundary, the pancreas, a kidney interface, and an ambiguous lesion, while skipping homogeneous liver parenchyma and a well-defined vessel. Unlike a post-hoc saliency map, these scores have an operational meaning: they determine whether native-resolution computation is executed.

## 7. Discussion

ErrorRoute treats computation as a resource that should be allocated according to expected segmentation benefit. This is distinct from simply making a backbone smaller. A uniformly lightweight model reduces representational capacity everywhere, whereas our coarse branch remains globally contextual and selectively recovers high-frequency detail where the expected payoff is high. The method is also distinct from uncertainty-guided refinement: uncertainty is useful evidence, but routing is supervised against realized improvement and therefore learns when uncertainty is actually actionable.

There are several limitations. First, utility targets require local refinement on sampled candidate blocks during training, increasing training cost even though inference is cheaper. Second, the current selector optimizes a hardware-agnostic proxy cost; direct latency-aware costs may be preferable on edge devices. Third, candidate blocks are axis-aligned cuboids, which can be inefficient for long curved structures. Future work could learn deformable candidate regions or combine routing with sparse 3D kernels.

## 8. Conclusion

We introduced ErrorRoute, a budgeted utility-aware routing framework for 3D medical image segmentation. The central idea is to predict the marginal benefit of native-resolution refinement and allocate compute to the blocks with the highest expected loss reduction per unit cost. A differentiable budgeted selector, local residual refiner, and cross-scale fusion produce a controllable accuracy– efficiency trade-off while preserving global anatomical context. Across diverse 3D segmentation benchmarks, utility-aware routing improves both computational efficiency and difficult-structure accuracy, suggesting a general path toward spatially adaptive medical vision systems.

## Impact Statement

Efficient 3D segmentation can reduce the compute and memory required for medical image analysis, potentially broadening access to volumetric models on constrained hardware. The method is not a diagnostic system and does not remove the need for dataset-specific validation, calibration, and clinical oversight. Efficiency gains should be evaluated alongside subgroup robustness and failure analysis before deployment.

## A. Notation

**Table 4.** Notation used in the paper.

| Symbol | Meaning |
| --- | --- |
| $x, y$ | input volume and segmentation |
| $z^c, z^f$ | coarse and final logits |
| $\mathcal{B}$ | multi-scale candidate block set |
| $c_i$ | compute cost of block $i$ |
| $\Delta_i$ | realized block loss reduction |
| $\nu_i$ | utility target per unit compute |
| $\hat{\nu}_i$ | predicted utility |
| $s_i$ | routing decision |
| $\rho$ | requested compute budget fraction |

## B. Additional Technical Details

### B.1. Candidate construction

At each training iteration we create three overlapping grids at native resolutions 32^3^, 48^3^, and 64^3^. To reduce candidate count, blocks containing less than 0.5% coarse foreground probability and low boundary gradient are subsampled during training but remain eligible at test time. All candidates share a common descriptor dimension after adaptive average pooling.

### B.2. Utility target estimation

For an expensive all-block oracle, the local refiner would be executed on every candidate. Instead, we sample up to 24 candidates per volume for target construction. Half are sampled proportionally to coarse error/boundary magnitude and half uniformly. The refinement loss used in Eq. 2 includes the block interior plus an 8-voxel context ring so that corrections that merely shift errors to the crop boundary are not rewarded.

### B.3. Budget solver

The training-time gate uses a straight-through Gumbel relaxation. The threshold *τ*_*B*_ is found by 12 iterations of bisection over the score range, which adds negligible overhead because the candidate set is small relative to the 3D backbone. Inference uses deterministic utility-per-cost sorting.

## C. Experimental Details

### C.1. Training protocol

All volumes are resampled to dataset-specific median spacing following the corresponding nnU-Net configuration. Intensity normalization uses z-score normalization for MRI and clipped z-score normalization for CT. We use random rotations, scaling, gamma augmentation, Gaussian noise, and mirroring. The coarse and local branches are optimized jointly; utility gradients do not propagate through the target construction path.

### C.2. Baseline configuration

Dense baselines are trained with the same patient splits and image preprocessing. SegBlocks-3D uses cubic blocks and three resolution levels. The uncertainty-refinement baseline selects blocks by predictive entropy at the same budget as ErrorRoute. The fixed coarse-to-fine baseline expands the coarse foreground bounding box by 24 voxels and refines the resulting crop at native resolution.

## D. Supplementary Results

**Table 5.** Routing strategy comparison at *ρ* = 0.3.

| Router | Dice | Oracle rec. | GFLOPs |
| --- | --- | --- | --- |
| Random blocks | 88.7 | 30% | 128 |
| Entropy | 89.8 | 62% | 128 |
| Boundary magnitude | 89.6 | 58% | 128 |
| Predicted error probability | 90.0 | 69% | 128 |
| Predicted utility (ours) | **90.5** | **83%** | 128 |
| Oracle utility | 91.5 | 100% | 128 |

**Table 6.** Cross-dataset router transfer. Utility estimators are trained on the source dataset and frozen on the target dataset.

| Transfer | Dice | Oracle recall |
| --- | --- | --- |
| AMOS22 → BTCV | 86.8 | 79% |
| BTCV → AMOS22 | 90.4 | 77% |
| KiTS19 → LiTS | 93.7 | 75% |
| Jointly trained target router | **94.2** | **83%** |

## E. Additional Discussion

The method can be viewed as a spatial analogue of conditional computation: instead of choosing which layer or expert to execute, ErrorRoute chooses which subvolumes deserve a higher-resolution computation path. The utility target provides a task-aligned credit signal for routing. Although we focus on segmentation, the same construction could apply to detection, registration, or quantitative measurement whenever a cheap global predictor and expensive local expert can be compared during training.

## References

Bilic, P., Christ, P. F., Vorontsov, E., Chlebus, G., Chen, H., Dou, Q., Fu, C.-T., Han, X., Heng, P.-A., Hesser, J., et al. The liver tumor segmentation benchmark (lits). Medical Image Analysis, 84:102680, 2023.

Hatamizadeh, A., Nath, V., Tang, Y., Yang, D., Roth, H., and Xu, D. Swin unetr: Swin transformers for semantic segmentation of brain tumors in mri images. In Brainlesion: Glioma, Multiple Sclerosis, Stroke and Traumatic Brain Injuries, pp. 272–284. Springer, 2022a.

Hatamizadeh, A., Tang, Y., Nath, V., Yang, D., Myronenko, A., Landman, B., Roth, H. R., and Xu, D. Unetr: Transformers for 3d medical image segmentation. In Proceedings of the IEEE/CVF Winter Conference on Applications of Computer Vision, pp. 574–584, 2022b.

Heller, N., Isensee, F., Maier-Hein, K. H., Hou, X., Xie, C., Li, F., Nan, Y., Mu, G., Lin, Z., Han, M., et al. The kits19 challenge data: 300 kidney tumor cases with clinical context, ct semantic segmentations, and surgical outcomes. arXiv preprint arXiv:1904.00445, 2019.

Isensee, F., Jaeger, P. F., Kohl, S. A., Petersen, J., and Maier-Hein, K. H. nnu-net: a self-configuring method for deep learning-based biomedical image segmentation. Nature Methods, 18(2):203–211, 2021.

Jang, E., Gu, S., and Poole, B. Categorical reparameterization with gumbel-softmax. In International Conference on Learning Representations, 2017.

Ji, Y., Bai, H., Ge, C., Yang, J., Zhu, Y., Zhang, R., Li, Z., Zhanng, L., Ma, W., Wan, X., et al. Amos: A large-scale abdominal multi-organ benchmark for versatile medical image segmentation. Advances in Neural Information Processing Systems Datasets and Benchmarks Track, 35, 2022.

Kendall, A. and Gal, Y. What uncertainties do we need in bayesian deep learning for computer vision? In Advances in Neural Information Processing Systems, volume 30, 2017.

Kervadec, H., Bouchtiba, J., Desrosiers, C., Granger, E., Dolz, J., and Ben Ayed, I. Boundary loss for highly unbalanced segmentation. In Proceedings of The 2nd International Conference on Medical Imaging with Deep Learning, volume 102, pp. 285–296. PMLR, 2019.

Litjens, G., Kooi, T., Bejnordi, B. E., Setio, A. A. A., Ciompi, F., Ghafoorian, M., van der Laak, J. A., van Ginneken, B., and Sánchez, C. I. A survey on deep learning in medical image analysis. Medical Image Analysis, 42:60–88, 2017.

Ma, J., Li, F., and Wang, B. U-mamba: Enhancing longrange dependency for biomedical image segmentation. arXiv preprint arXiv:2401.04722, 2024.

Milletari, F., Navab, N., and Ahmadi, S.-A. V-net: Fully convolutional neural networks for volumetric medical image segmentation. In 2016 Fourth International Conference on 3D Vision (3DV), pp. 565–571, 2016.

Roy, S., Koehler, G., Ulrich, C., Baumgartner, M., Petersen, J., Isensee, F., Jäger, P. F., and Maier-Hein, K. H. Mednext: Transformer-driven scaling of convnets for medical image segmentation. In Medical Image Computing and Computer Assisted Intervention – MICCAI 2023, pp. 405– 415. Springer, 2023.

Shazeer, N., Mirhoseini, A., Maziarz, K., Davis, A., Le, Q., Hinton, G., and Dean, J. Outrageously large neural networks: The sparsely-gated mixture-of-experts layer. In International Conference on Learning Representations, 2017.

Simpson, A. L., Antonelli, M., Bakas, S., Bilello, M., Farahani, K., van Ginneken, B., Kopp-Schneider, A., Landman, B. A., Litjens, G., Menze, B., et al. A large annotated medical image dataset for the development and evaluation of segmentation algorithms. arXiv preprint arXiv:1902.09063, 2019.

Verelst, T. and Tuytelaars, T. Segblocks: Block-based dynamic resolution networks for real-time segmentation. IEEE Transactions on Pattern Analysis and Machine Intelligence, 45(2):2400–2411, 2023.

Wang, X., Yu, F., Dou, Z.-Y., Darrell, T., and Gonzalez, J. E. Skipnet: Learning dynamic routing in convolutional networks. In Proceedings of the European Conference on Computer Vision, pp. 409–424, 2018.

Wu, Z., Nagarajan, T., Kumar, A., Rennie, S., Davis, L. S., Grauman, K., and Feris, R. Blockdrop: Dynamic inference paths in residual networks. In Proceedings of the IEEE Conference on Computer Vision and Pattern Recognition, pp. 8817–8826, 2018.

Xi, S., Wang, S., Safari, M., Hu, M., Tian, Z., and Yang, X. Uncertainty as a foundation for trustworthy medical imaging ai: A comprehensive review. 2025.

Xi, S., Hu, S., Wang, S., Ding, H., Li, Y., He, W., Zhang, K., Del Balzo, L., Zhong, C., Hu, M., et al. Grounding radiology report findings into medical image segmentation. npj Digital Medicine, 2026a.

Xi, S., Hu, S., Wang, S., Safari, M., del Balzo, L., Karim, E. U., Hu, M., Zhang, K., Wang, T., Weichselbaum, R. R., et al. A radiographic world model for clinical reasoning and evidence generation. arXiv preprint arXiv:2609.07719, 2026b.

